# Reduced dorsal CA1 Activity Limits Retention of the Temporal Component of Declarative Memory in the *Cntnap2* Knockout Mouse Model of Autism

**DOI:** 10.1101/2024.10.29.620866

**Authors:** Elise C. Rawlinson, Michael H. McCullough, Azza Sellami, Aline Marighetto, A. Shaam Al Abed, Nathalie Dehorter

## Abstract

Growing evidence implicates the hippocampus in the pathophysiology of autism spectrum disorder, particularly in the domains of social interactions and cognition. Yet, the mechanisms driving the hippocampal-dependent cognitive atypicalities in autism remain poorly defined. Here, we characterized how dysfunction of the CA1 subfield of the dorsal hippocampus drives abnormalities in critical components of declarative memory. Using trace fear conditioning in the *Cntnap2* knockout mouse model of autism, we found that capabilities to retain the association of temporally distant stimuli (i.e. temporal binding) were reduced relative to wildtype mice. Fiber photometry and optogenetic experiments demonstrated that reduced CA1 activity during temporal gaps underlies this impairment, linking CA1 dysfunction to a deficit in long-term memory retention in autism. Using a relational/declarative memory task, we also revealed a deficit in flexible spatial memory, and a preferential use of egocentric learning strategy. This inflexible learning strategy resulted from an imbalance between memory system activity, promoting frontostriatal-dependent procedural learning instead of dorsal CA1/hippocampus-dependent relational/declarative memory. Overall, this study establishes dorsal CA1 dysfunction as a circuit-level mechanism underlying cognitive inflexibility in autism, providing a neurobiological framework for hippocampal-dependent memory deficits in the condition.

## Introduction

Autism spectrum disorder (ASD) is a lifelong neurodevelopmental disorder comprising difficulties in social behavior, atypical communication skills, and the presence of restricted or repetitive behaviors or interests (DSM-5 and Association American Psychiatric, 2013). Beyond these core features, individuals with ASD present with difficulties in cognitive domains that can impact daily functioning and adaptability. In particular, people with ASD present with atypicalities in episodic/declarative memory (Bal et al., 2015), the ability to recall specific events and experiences (i.e., *what happened, when, and where*). However, the underlying neurological mechanisms of declarative memory impairments in ASD have yet to be fully elucidated.

While most research has focused on the dysregulation of the prefrontal cortex and the basal ganglia in ASD, recent studies have highlighted that hippocampal defects are key to the social domain of this phenotype (Banker et al., 2021). In addition, the hippocampus is known to play a critical role in declarative memory by forming temporally and spatially organized relational representations of past experiences, allowing flexible expression of memories (Eichenbaum, 2004). The CA1 subfield plays a key role in this relational process by sustaining the ability to link temporally distant events, known as temporal binding (Eichenbaum, 2014; Sellami et al., 2017). In autistic individuals, difficulties in the binding of episodic memories have been proposed in autistic individuals while memory for items alone remains relatively spared, positing a selective weakness in hippocampus-dependent linking of items and contexts (Bowler et al., 2011; Minor et al., 2023). In parallel with this relational hypothesis of memory defects, time estimation tasks in individuals with ASD have shown that disruptions in integrating temporal information affect the development of temporal relationships (Buonomano et al., 2023; Maister and Plaisted-Grant, 2011; Vogel et al., 2022). In this context, we hypothesized that CA1 activity underlying temporal processing and temporal linking is impaired in ASD, leading to declarative memory defects.

Here, we examined this hypothesis in the *Cntnap2* knockout (KO) mouse model of ASD, which exhibits autism-like phenotypes, including social behavior deficits (Peñagarikano et al., 2011; Peñagarikano and Geschwind, 2012), hippocampal alterations, and CA1 circuit-level changes and impairments in spatial discrimination (Paterno et al., 2021). Animals were assessed in two behavioral paradigms: trace fear conditioning (Pavlov, 1927; Sellami et al., 2017), a simple temporal binding-dependent memory task allowing the simultaneous recording and manipulation of CA1 activity; and a radial maze task modeling the complexity of temporal binding-dependent relational/declarative memory (R/DM). We demonstrate that *Cntnap2* KO mice present with mistuned dCA1 activation and abnormal reorganization of memory systems during temporal binding-dependent memory formation.

## Results

### *Cntnap2* KO mice demonstrate reduced temporal binding capability

We tested the temporal binding capabilities of the *Cntnap2* KO mouse model using the trace fear conditioning paradigm (TFC; **Figure 1A**), in which mice are required to memorize the association between a tone (Conditioned Stimulus) and a mild electric foot-shock (Unconditioned Stimulus), separated by a temporal gap. Successful conditioning across this gap is thought to depend on the organism maintaining a decaying memory representation, or *trace*, of the CS after its offset; accordingly, this gap is conventionally termed the trace interval, and the paradigm trace conditioning (Pavlov, 1927; Sellami et al., 2017). In the conditioning stage, independent groups of mice were exposed to three tone-shock pairings with a trace interval between 5 and 40 s (*i.e.* T5, T20, or T40; **Figure 1A**). All groups showed intact conditioning abilities, demonstrated by high and increasing freezing times (Blanchard et al., 1986) during the tones and traces periods (**Figure 1B, Figure S1A**). Twenty-four hours later, the animals were re-exposed to the conditioned *tone* in a neutral context (*i.e.* tone test) to assess 24h retention capacities of the tone-shock association. Retention was reflected in fear responses to the tone and trace compared with before and after the tone (*i.e.*, conditioned fear response in **Figure 1C**, % time freezing in **Figure S1B**). Elevated freezing during the pre-tone period at T40 may reflect fear generalization (**Figure S1B**, T20 vs. T40 (Sellami et al., 2017)). Interestingly, *Cntnap2* KO mice also displayed a strong and specific freezing response to the tone for up to a 20 s trace interval (**Figure 1C**, T20), but not for 40 s (**Figure 1C**, T40; **Figure S1B**, T40). The animals were then exposed to the conditioning *context* alone to evaluate contextual memory. As previously demonstrated (Al Abed et al., 2020b; Sellami et al., 2017), WT mice displayed efficient retention of the tone-shock association up to a 40-s trace interval, illustrated by a strong and specific conditioned fear response to the tone in a neutral context (**Figure 1D**). These results were replicated in both control and *Cntnap2* KO littermates (**Figure S1C**). Altogether, these results indicate that *Cntnap2* KO mice exhibit reduced temporal binding capacity compared to WT mice, which compromises the 24h retention of the tone-shock association.

**Figure 1.**
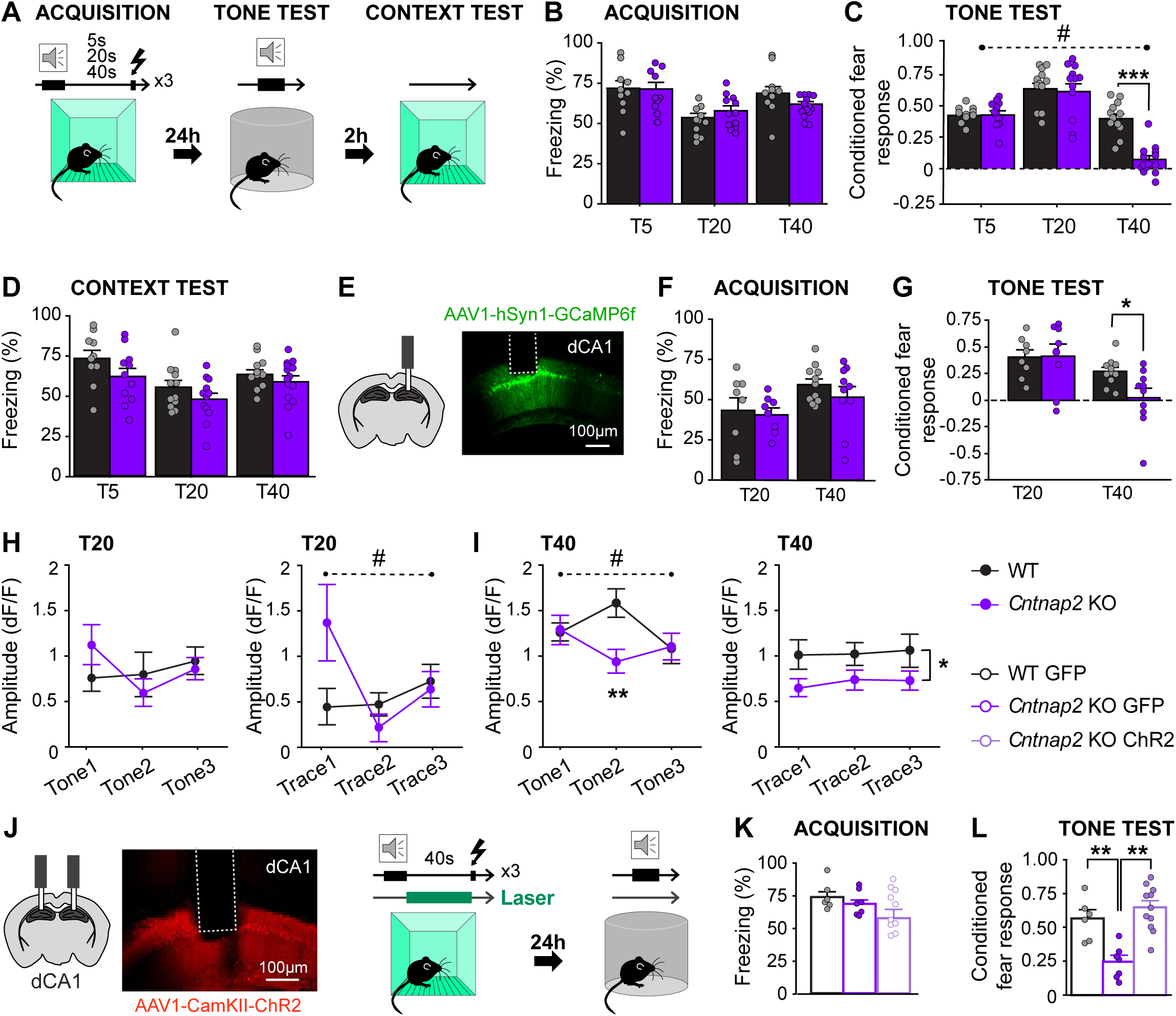
Reduced retention of temporal associations is associated with altered calcium responses during trace fear conditioning acquisition in *Cntnap2* KO mice. **A.** Experimental design: Trace Fear Conditioning includes a conditioning session in which mice have to associate a tone and a shock, separated by a time interval (trace). Independent groups of mice were trained with trace intervals of 5, 20, or 40 seconds. **B.** Freezing levels (percent) during the third tone of the conditioning session for WT (black) and *Cntnap2* KO (purple) groups, conditioned with distinct trace durations (T5: WT n=10; *Cntnap2* KO n=11; T20: WT n=12; *Cntnap2* KO n=11; T40: WT n=12; *Cntnap2* KO n=14). **C.** Conditioned fear response during the tone test (i.e., ratio of freezing before the tone compared to during the tone and the trace, see methods), 24h post-conditioning with distinct trace durations (Genotype X Trace interaction: F_2,65_=11.278; p<0.0001; T40: Genotype: F_2, 24_=6.944; p=0.0163). **D.** Freezing levels in the conditioning context (context test) for WT (black) and *Cntnap2* KO (purple) groups, 24h post-conditioning. **E. Fiber photometry. Left**: scheme depicting the fiber optic implantation targeting the dorsal CA1 subfield of the hippocampus (dCA1). **Right**: Representative image of the optic fiber placement, in the pyramidal cell layer of the dCA1, scale: 100µm. Calcium activity was recorded using the calcium indicator GCaMP6f (green). **F.** Freezing levels (percent) during the third tone of the conditioning session for WT (black) and *Cntnap2* KO (purple) groups, conditioned with distinct trace durations (T20: WT n=8; *Cntnap2* KO n=8; T40: WT n=11; *Cntnap2* KO n=10). **G.** Conditioned fear response during the tone test (i.e., ratio of freezing before the tone compared to during the tone and the trace, see methods), 24h post-conditioning with distinct trace durations (T40: Genotype: F_1, 19_=50.143; p<0.0001). **H. Left**: Peak amplitude of Ca^2+^ transients during the 3 tone presentations of the conditioning under a 20s trace-interval (ns). **Right**: Peak amplitude of Ca^2+^ transients during the 3 trace intervals of the conditioning under a 20s trace-interval (Genotype x Trace interaction: p=0.0476; WT n=8; KO n=8, trace 1: ns (p=0.0675)). **I. Left:** Peak amplitude of Ca^2+^ transients during the 3 tone presentations of the T40 conditioning session (Genotype x Tone interaction: p=0.0247; Tone: p=0.0053; WT n=11; KO n=10). **Right**: Peak amplitude of Ca^2+^ transients during the 3 trace intervals of the T40 conditioning session (Genotype: p=0.0408; WT n=11; KO n=10). **J. Optogenetics: Left** panel: scheme depicting the fiber optic implantations bilaterally targeting the dorsal CA1 subfield of the hippocampus (dCA1). **Right** panel: Representative image of the optic fiber placement, in the pyramidal cell layer of the dCA1, scale: 100µm. Pyramidal neurons activity was manipulated only during the trace interval using Channel Rhodopsin (ChR2). **K.** Freezing levels (percent) during the third tone of the conditioning session for GFP-injected WT (black, n=6), GFP-injected *Cntnap2* KO (purple, n=7), and ChR2-injected *Cntnap 2* KO (light purple/white, n=11) groups, conditioned with 40s trace duration. **L.** Conditioned fear response during the tone test (*i.e*., ratio of freezing before the tone compared to during the tone and the trace, see methods), 24h post-conditioning with distinct trace durations (Groups: F_2, 21_=15.576; p<0.0001; WT-GFP *vs* KO-GFP: p=0.0010; KO-GFP *vs* KO-ChR2: p<0.0001). Data shown at mean ± SEM. *: p<0.05; **: p<0.01; ***: p<0.001; * represents genotype effect. # represents genotype x variable interaction.

As freezing is the absence of movement, the well-reported hyperlocomotion in *Cntnap2* KO mice (Peñagarikano et al., 2011); **Figure S1D**) could represent a potential confounding factor. However, we observed comparable levels of resting time (**Figure S1D**, right panel) and similar freezing levels between WT and *Cntnap2* KO mice during acquisition, regardless of the trace interval (**Figure S1A**), as well as during memory tests under control conditions (**Figure S1B**; T5, T20, and T40). These findings indicate that locomotor activity alone cannot account for the observed differences in freezing. Taken together, our results demonstrate that *Cntnap2* KO mice exhibit reduced temporal binding capacity during conditioning, which does not affect acquisition but impairs the 24h retention of the tone-shock association.

### Mistuning of dorsal CA1 activity during temporal binding in *Cntnap2* KO mice

Whilst alterations in item binding in ASD have already been reported, the underlying mechanisms have not been described (Minor et al., 2023). As the sustained activity of dCA1 neurons during time intervals between stimuli is necessary for long-term retention of their association(Al Abed et al., 2020b; Sellami et al., 2017), we examined dCA1 activity during acquisition in *Cntnap2* KO mice. We recorded calcium transients during encoding and recall of trace fear conditioning, using fiber photometry in animals trained under a 20s or 40s-trace interval (**Figure 1E**). Both WT and *Cntnap2* KO mice implanted with an optic fiber could acquire the tone-shock association, reaching high levels of freezing after three tone-shock pairings (**Figure 1F, Figure S1E**). During conditioning, in both T20 and T40 conditions, regardless of genotype, the frequency of the calcium transients in response to the tone significantly decreased across tone presentations, suggesting neural adaptation to successive tone exposure (**Figure S1F, G**). *Cntnap2* KO mice trained under the 20s trace interval displayed no difference compared to WT in the frequency of calcium transients during the tone and the trace interval (**Figure S1F**), while the amplitude tended to be higher during the first trace interval in Cntnap2 KO mice (**Figure 1H**). In the 40s trace-conditioned group, the frequency of the calcium transients did not differ between WT and C*ntnap2* KO mice throughout conditioning (**Figure S1G**). In contrast, the calcium transient *amplitude* was significantly reduced both during the presentation of the second tone (**Figure 1I, left**) as well as throughout the three trace intervals in *Cntnap2* KO mice (**Figure 1I, right**). Overall, these results suggest a lack of synchronized neuronal activity during encoding, as previously reported (Simon and Wallace, 2016). During the tone test, as in Figure 1C, *Cntnap2* KO mice displayed appropriate retention of the tone-shock association when conditioned under a 20s-trace interval while the ones trained under a 40s-trace interval failed to express a fear response to the tone (**Figure S1H, I**). This indicates impaired retention of trace fear memory and temporal binding ability, relative to WT mice. The findings also suggest that sustained activity of dCA1 neurons during encoding of the three trace intervals, which is higher in WT than in *Cntnap2* KO mice, supports temporal binding and enables adequate retention of the association between the 40s-distant tone and shock despite their temporal separation without affecting initial acquisition. These results further support the idea that the reduction in temporal binding capacity in *Cntnap2* KO mice stems from a lack of sustained dCA1 activity during encoding and recall of the tone-shock association.

In WT mice, we have previously demonstrated that the activity of dCA1 principal neurons during the trace interval (but not outside of the trace interval) is required for long-term retention, but not for encoding of temporal associations (Sellami et al., 2017). To further confirm a causal link between sustained dCA1 activity during temporal gaps and 24h retention in *Cntnap2 KO*, we optogenetically activated principal neurons specifically during the 40s trace interval of the conditioning session using ChannelRhodopsin (ChR2; **Figure 1J**), while leaving neuronal activity unaltered during the remainder of acquisition and the 24h test. During the conditioning session, all groups displayed increased freezing across the three tone-shock pairings, indicative of efficient acquisition (**Figure 1K, Figure S1J**). Twenty-four hours later, ChR2-expressing *Cntnap2* KO mice demonstrated a specific freezing response to the tone and trace, unlike GFP-expressing *Cntnap2* KO mice (**Figure 1L, Figure S1K**). These results demonstrate that dCA1 neuronal activity during trace intervals drives temporal binding in *Cntnap2* KO mice.

### Egocentric learning and inflexible long-term memory in *Cntnap2* KO mice

Following the observation that *Cntnap2* KO mice present reduced dCA1 activity associated with temporal binding capabilities, we examined potential atypicalities in declarative memory. We used a radial maze task that models the complexity of temporal binding-dependent relational/declarative memory (R/DM) (Sellami et al., 2018). This task assesses the characteristic flexibility of R/DM expression, which relies on the dCA1-dependent ability to relate temporally distant events (Sellami et al., 2017).

To do so, the R/DM task is divided into two stages: an acquisition phase and a flexibility test (**Figure 2A**). In the acquisition stage, mice are trained daily to learn the food-reward location within 3 separate pairs of adjacent arms. Critically, this task structure requires mice to encode each pair as a distinct, temporally separated experience, rather than allowing continuous comparison between pairs. Cumulative evidence shows that declarative flexibility relies on a hippocampus-dependent relational representation of separate experiences formed for each pair during the acquisition phase (Al Abed et al., 2020b; Mingaud et al., 2007; Sellami et al., 2017). After reaching the learning criterion (>75% correct choices of the rewarded arm), declarative memory is then probed by testing whether animals can recombine two of these separately learned pairs (A and B) into a novel configuration (pair AB) without any change to the reward location **(Figure 2A**). Successful performance requires that the two pairs were encoded relationally, within a shared spatial representation, allowing this new relation to be inferred; poor performance, in contrast, suggests animals relied on rigid, pair-specific response rules that do not generalize to novel arm combinations. To impose a temporal binding demand on relational memory acquisition, the presentation of each pair is separated by a 20s inter-trial interval. This timing was empirically determined in (Sellami et al., 2018, 2017).

**Figure 2.**
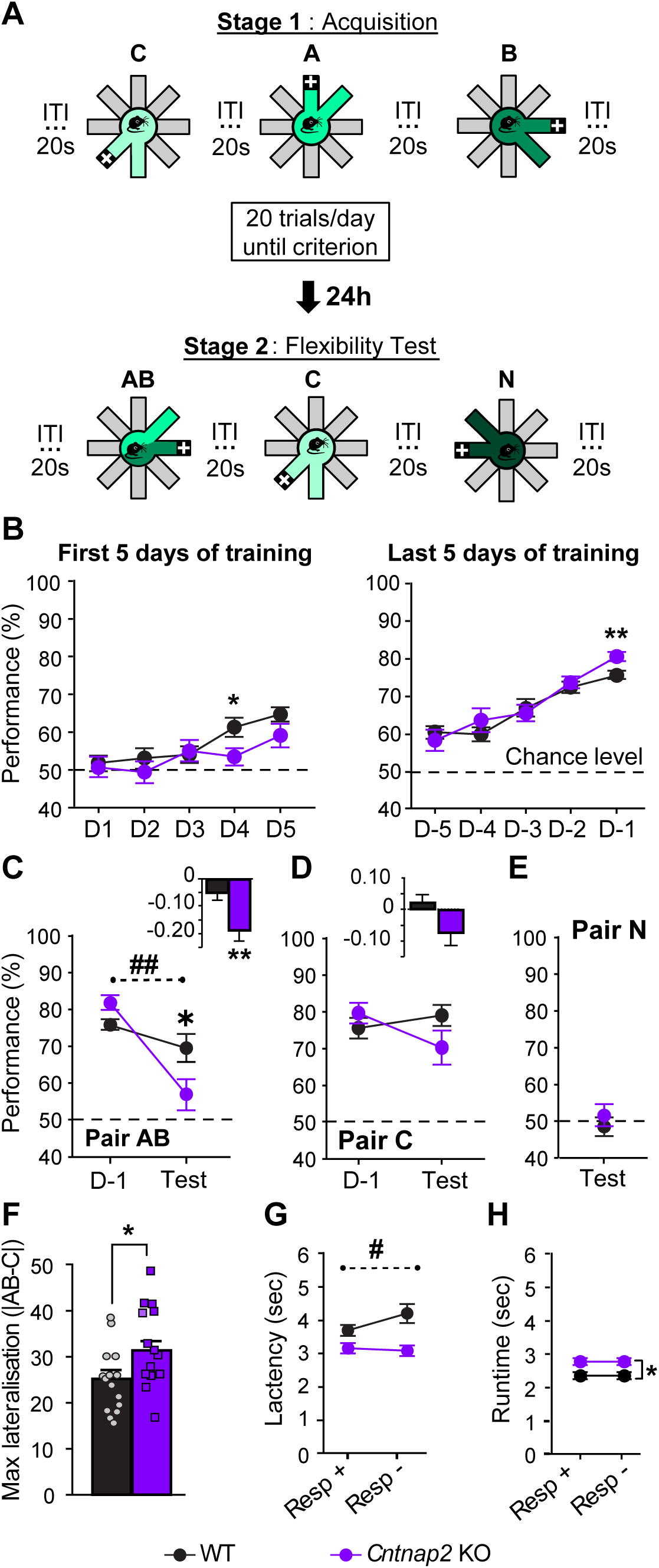
*Cntnap2* KO mice display impaired relational/declarative memory. **A.** Experimental design: Relational/Declarative memory task. ITI: Inter-trial interval; +: represent reward location within each pair of arms. **B.** Average correct answers for WT (black; n = 15) and *Cntnap2* KO mice (purple; n = 14) for pairs A, B, and C during the 5 first days of acquisition (left panel; day (D); D4: Genotype: p=0.0149), and the last 5 days of acquisition (right panel; D-5: 5 days before they reach learning criterion; D-1: Genotype: p=0.0082). **C.** Comparison of the correct answers during the last day of training (D-1) averaged on pairs A and B, and during the flexibility test on the recombined pair AB (Genotype X Performance interaction: F_1,27_=12.639, p=0.0014; Test, Genotype: p=0.0239). Inset: D-1 *vs.* Test performance index on pairs AB (Genotype: p=0.0022). **D.** Comparison of the correct answers on pair C during the last day of training (D-1) and the flexibility test. Inset: D-1 *vs*. Test performance index for Pair C. **E.** Average performance on the new pair N during the flexibility test. **F.** Maximum lateralization index during stage 1 (absolute value of the average performance between pairs A and B (left baited) minus Pair C (right baited); Genotype: p=0.0153). **G.** Average decision latencies for correct (+) and incorrect (-) answers throughout training (Genotype X Response interaction: F_1,31_=7.960, p=0.0083). **H.** Average runtimes for correct (+) and incorrect (-) answers throughout training (Resp -, Genotype: p=0.0500). Dashed line illustrates chance level. Data shown at mean ± SEM. *: p<0.05; **: p<0.01; # shows genotype x performance interaction.

Both wild-type (WT) and *Cntnap2* KO mice were able to acquire the task (**Figure 2B**). *Cntnap2* KO mice tended to require more days to reach criterion (average number of days: WT: 8.06, KO: 10.86, ns, raw data in Figure S3) and reached higher performance levels at the end of training (**Figure 2B**, right panel).

In the flexibility test, their performance dropped significantly between the last training session and the flexibility test (stage 2 of the R/DM task), and *Cntnap2* KO mice performed significantly worse than WT (**Figure 2C** - Recombined pair AB). In contrast, both groups performed similarly on the control pairs: the unchanged pair C, with high performance **(Figure 2D)**, and the novel pair N, at chance levels **(Figure 2E).**

What could explain this lack of flexibility? We have previously shown that the flexibility component of R/DM depends on the learning strategy used during the acquisition stage: either allocentric spatial mapping (relational) or egocentric (procedural), the latter leading to lateralization. Indeed, when the use of spatial cues is impaired, as in cases of hippocampal damage or dysfunction, mice adopt a left-right egocentric strategy to memorize reward locations leading to inflexible memory (Mingaud et al., 2007; Sellami et al., 2017). Therefore, the chance-level flexibility performance of *Cntnap2* KO mice suggests a bias towards a procedural learning strategy, in line with the well-documented prominence of behavioral inflexibility (Uddin et al., 2015) and the use of egocentric learning strategies in ASD (Brunner et al., 2015; Ring et al., 2018). This was confirmed by analyzing lateralization during acquisition by comparing performance on left-baited pairs (pairs A and B), *vs.* right-baited pair C (Methods, **Figure 2F**). *Cntnap2* KO mice displayed significantly more lateralization than WT, meaning they used an egocentric/procedural strategy to learn the position of the food reward rather than using the spatial mapping strategy observed in WT mice. We found no significant correlation between the use of egocentric strategy and the number of days to reach criterion (Data in Figure S3). Notably, performance during Day 4 of acquisition in WT mice was significantly negatively correlated with their use of lateralized strategy (r^2^ = -0.518, p=0.0471), which was not seen in *Cntnap2* KO mice. Together, this further confirms that *Cntnap2* KO mice employ a lateralized strategy to learn the task, which impacts both task performance (**Figure 2B**, day 4), and memory flexibility (**Figure 2C**).

Interestingly, *Cntnap2* KO mice had shorter decision times than WT **(Figure 2G)**, consistent with the well-demonstrated impulsivity observed in ASD (DSM-5 and Association American Psychiatric, 2013). In addition, WT mice demonstrated a longer time to enter a non-baited arm than a baited arm, suggesting that they evaluate reward location before entering the arm, likely relying on deliberation and precise spatial representation. In contrast, *Cntnap2* KO mice showed similarly short latencies regardless of their choice, suggesting impulsive, automatic responses and impaired spatial representation of reward information. In addition, once a choice was made, their running time (from the central platform to the end of the arm) was longer than that of WT mice (**Figure 2H**), implying that while the deliberative phase is shortened or absent in KO mice, there may be a compensatory adjustment following impulsive decisions.

Altogether, these findings indicate that *Cntnap2* KO mice exhibit impaired flexible memory expression, consistent with a relational/declarative memory deficit and increased reliance on an egocentric/procedural learning strategy.

### Potential shift towards striatum-centered activities in *Cntnap2* KO mice

Each learning strategy is known to depend on partially distinct brain systems: hippocampus-dependent systems supporting allocentric spatial/relational learning, and striatum-based systems leading to egocentric/procedural learning (White and McDonald, 2002). To assess learning-induced activity in these different structures, we quantified the levels of the cellular activity marker cFos after day 3 of R/DM task acquisition in structures involved in memory (i.e., the dorsal hippocampus [dCA1, dCA2, dCA3, the dorsal dentate gyrus (dDG)]; the dorso-medial (DMS) and the dorso-lateral striatum (DLS); the prefrontal cortex [infralimbic (IL), prelimbic (PrL), and anterior cingular (ACC)]; **Figure 3A, Figure S2**).

**Figure 3.**
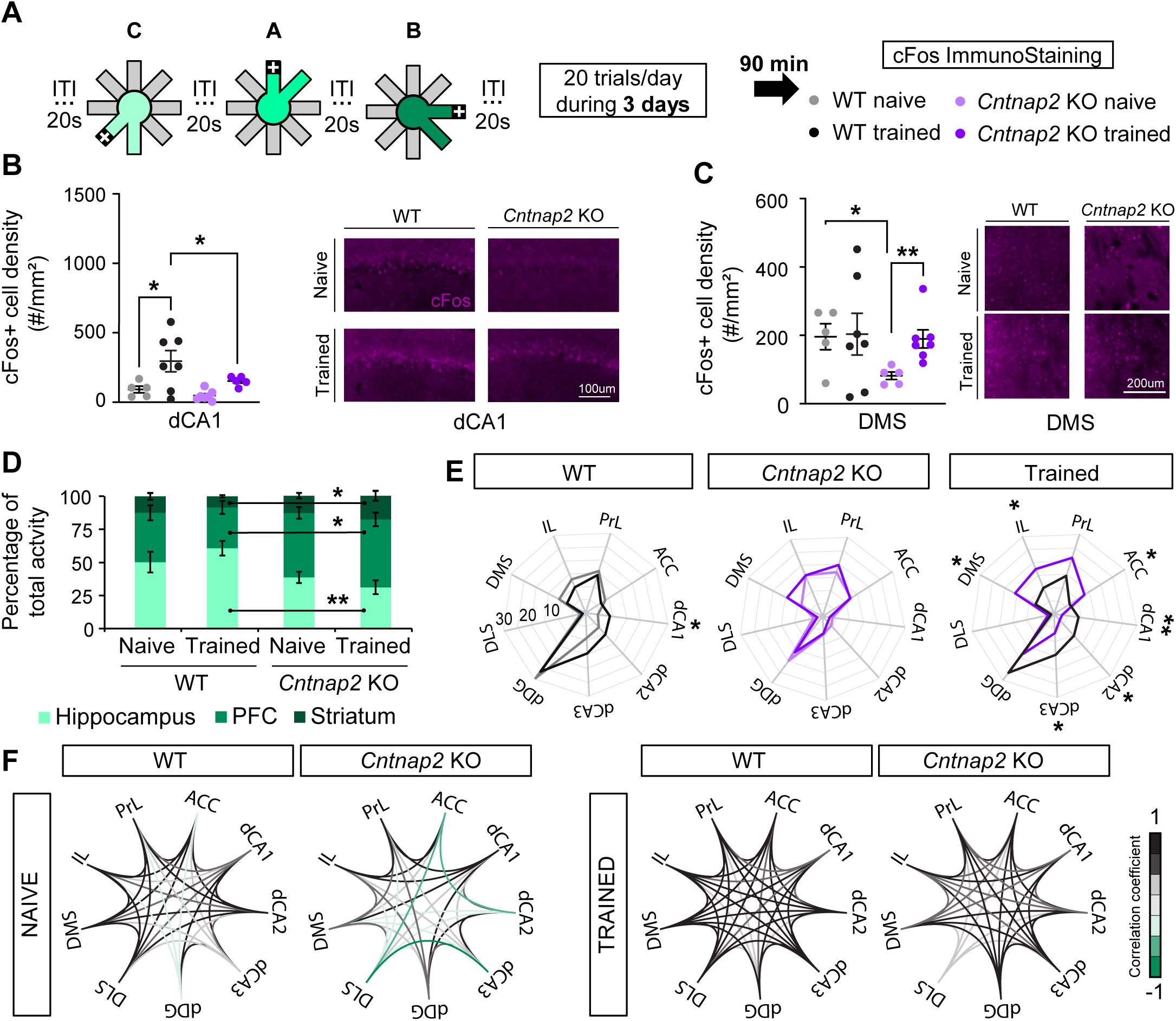
Learning-induced activity and functional connectivity in *Cntnap2* KO mice. **A.** Experimental design: cFos levels were quantified 90min after the 3^rd^ training session of the Relational/Declarative memory task. ITI: Inter-trial interval; + represents reward location within each pair of arms (WT Naïve: n=5; WT Trained: n=7; *Cntnap2* KO: n=5; *Cntnap2* KO Trained: n=7). **B.** number of cFos-positive neurons (cell density/mm^2^) in the dorsal dCA1 (WT naïve *vs* Trained: p=0.0491; Trained WT vs. KO: p=0.0129) with representative images of the cFos staining (Magenta) in dCA1, scale: 100µm, for naïve (light colors) and trained (dark colors) WT (grey/black) and *Cntnap2* KO (purple) groups. **C.** number of cFos-positive neurons (cell density/mm^2^) in the Dorso-Medial Striatum (DMS; Naive WT vs. KO: p=0.0189; KO Naïve vs Trained: p=0.0085) with representative images of the cFos staining (Magenta) in DMS, scale: 200µm. **D.** Percentage of total activity: activity of each structure over to the total activity measured (percentage; Trained WT vs. KO: Hippocampus: p=0.00194; PFC: p=0.00107; Striatum: p=0.0382). **E.** Percentage of total activity: activity each subfield within the total network, in WT mice (naïve vs trained, left panel; dCA1: p=0.0242), in *Cntnap2* KO mice (Naïve vs trained, middle panel), and in trained mice (WT vs *Cntnap2* KO mice, dCA1: p=0.009; dCA2: p=0.0174; dCA3: p=0.0274; ACC: p=0.0438; DMS: p=0.0350; IL: p=0.0267; PrL: p=0.0476; right panel, same data as in left and middle panels). **F.** Pairwise between-structure correlation strength of cFos cell density. Data shown as mean ± SEM. *: p<0.05; **: p<0.01, ***: p<0.001.

As previously shown (Al Abed et al., 2020b; Sellami et al., 2017), we found a significant activation of dCA1 in trained WT mice compared to naïve WT mice (**Figure 3B**), consistent with the use of an allocentric/spatial learning strategy (White and McDonald, 2002). In contrast, *Cntnap2* KO mice showed no learning-induced dCA1 activation (**Figure 3B**), together with hypoactivation of dCA3 in both naïve and trained groups compared to WT mice (**Figure S2A**). This reduced hippocampal activation may contribute to the observed deficit in temporal binding and spatial learning capabilities in this task. Moreover, *Cntnap2* KO mice displayed a significant training-induced activation of the DMS compared to their naïve controls, a pattern not observed in WT mice (**Figure 3C**). No other significant changes were detected in the hippocampus (*i.e.* dCA2, DG; **Figure S2A**), in the DLS (**Figure S2B**) or in the prefrontal cortex regions (*i.e.*IL, PrL, ACC; **Figure S2C**). Together, these findings suggest increased engagement of DMS-associated networks during the egocentric learning strategy adopted by *Cntnap2* KO mice.

To further dissect the balance between memory systems, we analyzed the relative contribution of each structure within the total activation of the system (**Figure 3D-E**). After training, we found a disproportionate representation of the PFC and striatum relative to the hippocampus in *Cntnap2* KO mice compared to WT (**Figure 3D**). Indeed, compared to WT naïve mice, WT trained mice exhibited an increase in the weight of the dCA1 in the network activity, likely supporting temporal binding and, through ensuing relational organization of the 20s-distant pair presentations, the formation of flexible memory. In contrast, no such reorganization of network activity was observed in *Cntnap2* KO mice with training (**Figure 3E**). We observed differences in the relative contribution of individual substructures between trained WT and *Cntnap2* KO mice. Specifically, training was associated with greater involvement of the dCA1, dCA2, and dCA3 in WT mice, whereas *Cntnap2* KO mice showed greater involvement of the DMS, IL, and ACC. Overall, we observed compensatory activation of the frontal-striatal structures concomitant with hippocampal hypoactivation in the *Cntnap2* KO model.

In addition to the learning-induced activity of individual structures, pairwise correlation of activity between structures enhances our understanding of network-wide synchronization. While naïve WT mice exhibited mainly positive correlations between the striatum, PFC, and hippocampus, with the exception of the dDG which was negatively correlated with the PFC, naïve *Cntnap2* KO mice displayed predominantly negative correlations (**Figure 3F**).

This indicates that, at baseline, inter-structural relationships are dysregulated in *Cntnap2* KO mice, potentially limiting the recruitment of a hippocampus-based learning strategy. Altogether, the analysis of learning-induced brain activity suggests a shift towards striatum-centered activity and connectivity in *Cntnap2* KO mice, likely promoting an egocentric learning strategy and resulting in inflexible memory.

## Discussion

This study establishes a novel mechanistic framework that builds on prior reports of hippocampal alterations in *Cntnap2* KO mice (Paterno et al., 2021). It demonstrates that compromised CA1 hippocampal activity during encoding disrupts temporal binding, which, by impairing the relational processing of temporally distant events, leads to deficits in retention of declarative memory. Moreover, hippocampal dysfunction in *Cntnap2* KO mice appears to shift learning away from hippocampus-dependent strategies and toward egocentric strategies.

### Interplay between hippocampus and striatum in memory deficits in *Cntnap2* KO mice

Individuals with ASD show atypicalities in declarative memory, characterized by impaired relational binding, i.e., the ability to link discrete items or events into a unified representation, while memory for individual items in isolation appears comparatively preserved (Minor et al., 2023). Because many real-world relations must be formed across a temporal gap (e.g., linking an event to its outcome, or one experience to another encountered later), we propose that this relational impairment may arise, at least in part, from a deficit in temporal binding: the hippocampus-dependent process of maintaining and integrating information across time to construct coherent, flexible memories.

Here we demonstrate a reduction in dCA1-dependent temporal binding capacity in *Cntnap2* KO mice, which could contribute to the declarative memory impairment by disrupting the relational organization of temporally distant events. Indeed, in the trace fear conditioning experiments *Cntnap2* KO mice showed efficient temporal binding with time intervals up to 20s, compared with 40s in WT mice, and this reduction was associated with a reduction of dCA1 activity during the encoding of the 40s-trace fear conditioning. Temporal integration of stimuli relies on hippocampal activity, in particular that of dCA1 (Buzsáki and Tingley, 2018; Chowdhury et al., 2005; Sellami et al., 2017; Tsien et al., 1996). Similarly, we found an impaired activation of the dCA1 in *Cntnap2* KO mice during the acquisition of the Relational/Declarative Memory (R/DM) task (**Figure 3B**).

The lack of training-induced dCA1 activation likely prevents fine-tuned temporal integration of distant events, leading to flexibility deficits (Chang and Gold, 2003; Etchamendy et al., 2003; Mingaud et al., 2007; Sellami et al., 2017). Moreover, analysis of broader memory systems’ activity during encoding revealed an aberrant recruitment of the DMS in the *Cntnap2* KO mice, along with an altered pair-wise correlation between activation of related brain regions (**Figure 3F**). While WT mice exhibited predominantly positive correlations, *Cntnap2* KO mice showed negative correlations among the DLS, prefrontal cortex and hippocampus. This pattern was also observed in naïve groups, suggesting that the bias in ASD towards lateralized learning strategy stems from baseline mistuning of functional connectivity between brain structures necessary for spatial learning in ASD.

This shift toward a striatum-centered system in ASD also implies a predominance of unconscious encoding, in which relations between items are acquired via stimulus–response associations, leading to inflexible, habit-like actions (Ferbinteanu, 2016). Whether the increased reliance on corticostriatal pathways reflects a compensatory adaptation to impaired hippocampal processing or a direct consequence of *Cntnap2* deficiency remains an open question. *Cntnap2* is broadly expressed across cortico-hippocampal and corticostriatal circuits and has been implicated in neuronal connectivity, synaptic function, and network synchronization (Lazaro et al., 2019; Scott et al., 2019; Selimbeyoglu et al., 2017). It is therefore possible that the altered engagement of corticostriatal networks observed here is not solely secondary to reduced hippocampal activity, but also reflects primary circuit alterations caused by the genetic mutation. Future studies directly examining corticostriatal activity and manipulating these circuits during learning will be required to disentangle primary genetic effects from compensatory network adaptations. However, additional mechanisms may contribute to the phenotype. *Cntnap2* KO mice exhibit other neural and behavioral alterations (Peñagarikano et al., 2011), including epileptic seizures and hyperlocomotion, suggesting that CA1 hypoactivity is likely only one of several factors influencing performance. Similarly, although previous work demonstrated that dCA1 activation rescues temporal binding only when delivered during the trace interval (Sellami et al., 2017), future studies should determine whether the same temporal specificity is maintained in the *Cntnap2* KO model.

### Decoding spatial and temporal maps in the *Cntnap2* KO mouse model of autism

The CA1 region is recruited with other hippocampal areas to form and maintain spatial, temporal and social maps, which are mental relational representations of our environment (Barrientos and Tiznado, 2016). CA1 neurons contribute to encoding and recalling spatial information, enabling individuals to navigate and understand their surroundings. Specific “time cells” create a temporal framework that supports episodic memory, encoding the passage of time and the prediction of future events (Eichenbaum, 2014). Recent research also suggests that CA1 contributes to social cognition through so-called “social maps” which process and recall social information that is essential for effective social behavior and social dynamics (Zhang et al., 2022). Yet, little is known about how cognitive deficits evolve over time and their long-term effects in ASD. While the present study focused on dCA1 activation during acquisition to assess how dCA1 binds distinct stimuli into unified representations, future investigations directly examining the causal role of dCA1 activity in memory maintenance and retrieval (*e.g.* ArchT-mediated silencing during recall) will be essential for a comprehensive understanding of dCA1’s role in memory processes in ASD, as well as for exploring how specific maps are formed and maintained in this condition.

The role of dCA1 in spatial representations has been profusely documented. Consistently, we recently demonstrated the central role of dCA1 in the maladaptive contextual encoding in post-traumatic stress disorder (PTSD), i.e., the degradation of declarative memory surrounding a highly stressful event (Al Abed et al., 2020a; Brewin, 2011; DSM-5 and Association American Psychiatric, 2013). Interestingly, growing evidence reports the propensity to develop PTSD in ASD (Al Abed et al., 2024; Rumball et al., 2020), suggesting that reduced dCA1 activation may impact cognition beyond everyday learning. Together, these studies further underscore the hippocampus as a promising target to alleviate ASD pathophysiology.

To obtain a deeper understanding of the hippocampus computation underlying memory formation, we analyzed the correlation between neuronal dynamics and behavioral outcomes within the fear memory test. Fiber photometry recordings in the dCA1 showed that whilst WT mice displayed sustained hippocampal activity during trace interval (**Figure 1I**), hippocampus-dependent memory systems were not similarly activated in the *Cntnap2* KO mice (**Figure 3B, E**). We also found a marked decrease in the amplitude of calcium transients in *Cntnap2* KO mice (**Figure 1I**). This profile was not observed when *Cntnap2* KO mice were trained within their capacity (*i.e.* 20 s trace interval; **Figure 1H**). However, because fiber photometry provides a population-level measure of activity, it does not resolve the contribution of individual CA1 neurons or underlying cellular and synaptic mechanisms. Future studies using approaches with single-cell or subcellular resolution will be required to determine how these mechanisms contribute to the altered CA1 dynamics observed here.

Overall, these results suggest that the deficits in temporal binding and declarative memory originate from reduced hippocampal neuronal synchronization and dynamics. An additional possibility is that sustained CA1 activity during the trace interval is supported not only by recurrent network interactions but also by intracellular mechanisms within CA1 pyramidal neurons. Persistent dendritic calcium signaling and CaMKII-dependent processes, including dendritic, delayed and stochastic CaMKII activation (Jain et al., 2024) and short-term associative plasticity of calcium dynamics (Caya-Bissonnette et al., 2023; Caya-Bissonnette and Béïque, 2024), have been proposed to support prolonged information retention. Determining the relative contributions of these intracellular and circuit-level processes will require future investigations with single-cell and subcellular resolution, as the network-level approaches used in the present study did not allow us to distinguish whether the sustained activity observed here arises from these cellular mechanisms, recurrent circuit dynamics, or a combination of both. In the same line, a recent study reported that neurons allocated to the memory trace displayed freezing-specific activity compared to non-allocated neurons during contextual fear conditioning (Mocle et al., 2024). While our study design does not provide these details, the lack of active cells in the dCA1 during acquisition of the relational/declarative memory task could imply memory trace allocation to the striatum rather than the hippocampus (Rashid et al., 2016). Further experiments are needed to determine whether atypical engram allocation is linked to learning strategies in ASD.

### Dissociating hyperactivity from impulsivity in *Cntnap2* KO mice

As *Cntnap2* KO mice display well-documented hyperlocomotion (Peñagarikano et al., 2011; **Figure S1D**), a general increase in motor output represents a potential confound for several of the behavioral readouts in this study. We addressed this directly for the TFC paradigm, where resting time did not differ between genotypes despite increased total distance and velocity in the open field (**Figure S1D**), and where freezing levels during acquisition were comparable across trace intervals and genotypes (**Figure S1A**). These findings indicate that the genotype-specific loss of retention at the 40 s trace interval reflects a deficit in temporal binding rather than an inability to freeze.

However, in the R/DM task our data more directly implicate impulsivity, rather than hyperactivity, as a contributing factor to task performance. *Cntnap2* KO mice showed shorter decision latencies specifically at the choice point of the maze (**Figure 2G**), and, critically, this latency reduction was similar regardless of whether the eventual choice was correct or incorrect. In contrast, WT mice took longer to enter non-baited arms, consistent with deliberation over reward location before committing to a choice. This pattern is consistent with premature, automatic responding rather than a general increase in locomotor drive: KO mice were in fact slower once a choice had been made (*i.e.* longer runtimes on both incorrect and correct trials; **Figure 2H**), arguing against a simple speed-accuracy effect driven by hyperactivity.

This distinction is conceptually important, as impulsivity, characterized by premature responding and reduced response control, is a well-documented feature of ASD in both clinical and preclinical settings (DSM-5 and Association American Psychiatric, 2013), including in other autism-relevant mouse models assessed with operant impulse-control paradigms. Examples include the 5-choice serial reaction time task (CSRTT), where increased premature responding has been reported independently of general locomotor activity (McTighe et al., 2013). We therefore interpret the shortened decision latencies in *Cntnap2* KO mice as reflecting reduced deliberative evaluation of spatial/reward information prior to responding, which may compound, but is dissociable from, the reduced hippocampal engagement and egocentric strategy bias described above.

We note, however, that our task was not designed to formally quantify impulsivity (*e.g.,* via premature responses during an imposed delay, as in the 5-CSRTT), and future studies directly assessing response inhibition in *Cntnap2* KO mice will be required to determine whether this impulsive-like phenotype is mechanistically linked to, or independent of, dCA1 hypofunction.

### Aged-like performance in *Cntnap2* KO mice suggests early cognitive decline

In this study, we demonstrate the disruption in hippocampal and CA1 functioning in the *Cntnap2* KO mice, a hallmark of memory defects observed during aging (Etchamendy et al., 2012; Leal and Yassa, 2015). Consistent with reports suggesting that people with autism are at risk of premature cognitive aging (Wang et al., 2024), we have found that young adult *Cntnap2* KO mice exhibit a memory profile similar to those of aged WT mice (Al Abed et al., 2020b; Marighetto et al., 2011; Sellami et al., 2017). Compared to young WT mice, both groups showed flexibility impairments in the relational/declarative radial maze task, and both exhibited a striatum-biased learning strategy. However, the underlying mechanisms driving these deficits remain unknown.

We previously demonstrated that dCA1 activity during the temporal gaps is necessary to link separate pairs into a relational/flexible memory (Al Abed et al., 2020b; Sellami et al., 2017) and that targeted dCA1 optogenetic activation during these intervals increases temporal binding capacity, rescuing relational/declarative memory formation. Given that CA1 neurons are less synchronized in ASD, targeted dCA1 activation could alleviate memory deficits. We propose that fine-tuning dCA1 activity may normalize memory system balance, offering a novel therapeutic approach for cognitive impairments in ASD. Further delineation of associated connectomes (Hollunder et al., 2024) could provide insights into targeted brain circuit therapies.

Autism is a highly heterogeneous condition, with individuals presenting a wide range of atypical cognitive abilities. Consequently, the extent and nature of memory deficits vary considerably across the spectrum. Research has yet to fully elucidate how these deficits manifest, particularly in relation to other cognitive domains such as attention, executive functioning, and language. Sex differences are another factor to consider (Van Wijngaarden-Cremers et al., 2014). While this study included both male and female mice, we did not observe sex-specific effects, suggesting that the behavioral and cognitive phenotypes examined in the *Cntnap2* KO mouse model are largely independent of sex. This is consistent with previous studies reporting no robust sex differences (Al Abed et al., 2024).

A better understanding of these individual differences is necessary to create more personalized and effective interventions. As such, further studies will have to assess these aspects in different models to fully comprehend the common cognitive mechanisms at play in autism.

## Methods and material

### Mice

3 to 5-month-old naive mice were collectively housed in standard Makrolon cages, in a temperature- and humidity-controlled room under a 12h light/dark cycle (lights on at 07:00), with *ad libitum* access to food and water. We used both males and females to minimize the number of mice produced in this study. *Cntnap2*^+/+^ (WT) and *Cntnap2*^-/-^ (KO) mice (Jackson ID: #017482 (Peñagarikano et al., 2011)) were non-littermate, though both lines were originally derived from breeders of the same colony and were used for behavior and histological analyses. To minimize potential differences between strains, we regularly inter-crossed the wildtype and *Cntnap2* KO colonies (*i.e.*, every year) to generate heterozygous mice. These heterozygotes were then bred to produce either WT or KO mice for experiments. Throughout the study, we used non-littermate mice originating from the same breeders. To ensure that observed differences between WT and *Cntnap2* KO were not influenced by this breeding strategy, we also assessed trace fear conditioning capacities in littermate mice and confirmed our findings using Het × Het breeding experiments (**Figure S1**).

All mice were bred in the same room, and maintained under identical husbandry and rearing conditions, including cage content, bedding, diet, and light/dark cycle. All experiments took place during the light phase. We replicated the behavioral experiments in two to four different batches. Every effort was made to minimize the number of animals used and their suffering. All procedures were conducted in accordance with the European Directive for the care and use of laboratory animals (2010-63-EU) and the animals care guidelines issued by the animal experimental committee of Bordeaux University (CCEA50, agreement number A33-063-099; authorization N°21248), and from the Australian National University Animal Experimentation Ethics Committee (protocol numbers A2018/66, A2020/26, A2021/43, and A2024/362).

### Trace fear conditioning task (TFC)

#### Apparatus

Fear conditioning behavior was performed in a Plexiglas conditioning chamber (20 x 20 x 30 cm), in a brightness of 100 lux, given access to the different visual-spatial cues in the experimental room. The floor of the chamber consisted of stainless-steel rods connected to a shock generator (Imetronic, Pessac, France).

#### Trace fear conditioning procedure

The box was cleaned with 70% ethanol before each trial. *During conditioning*, each animal received 3 pairings of a tone (70 dB, 1 kHz, 30 s) and a footshock (0.3 mA, 50 Hz, 1 s). The two stimuli were separated by a trace interval of 5, 20, or 40 s, depending on the group. Each tone-shock pairing was separated by a 1min-interval.

All mice were submitted 24 hours later to the *Tone retention test* during which they were re-exposed to the tone alone in a dark and modified chamber [circular opaque white chamber cleaned with a 1% acetic acid solution; 2min pre-tone, 2min tone and 2min post-tone]. A context test was also performed 2 hours later, during which mice were re-exposed to the conditioning chamber alone for 6 min.

Animals were continuously recorded for off-line scoring of freezing, by an experimenter ignorant of the experimental condition/group. Tone test was scored by 2 independent scorers. Freezing is defined as a lack of all movement except for respiratory-related movements. Acquisition of the TFC was measured by the progression of freezing during the tone across the 3 tone deliveries. 24h-retention of the trace association was measured during the tone test by the evolution of percentage of freezing time. Conditioned fear response to the tone was evaluated by calculating a normalized ratio between the tone and trace periods, *versus* outside these periods (i.e., Pre-Tone and Post-Tone): [(Tone+Trace)/2 - (pre-Tone+post-Tone)/2] / [(Tone+Trace)/2 + (pre-Tone+post-Tone)/2].

### Radial-maze task of Relational/Declarative memory (R/DM)

#### Apparatus

We used an open 8-arm radial maze made of grey Plexiglas, automatized by videotracking (IMETRONIC – Pessac, France). The diameter of the central platform is 30 cm, and the arms are 55 cm long by 10 cm large. Each arm is equipped with a door at its entrance and a food-pellet delivery system at its end. The doors are individually controlled (raised up or dropped down) by the computerized system which also controls pellets availability in the food tray at the end of each arm individually, according to the task. The maze is placed in an empty room containing visual cues to enable spatial discrimination.

#### Behavioral Procedure

During the entire procedure, animals were submitted to one daily session. Prior to memory testing, animals were habituated to the apparatus over a period of two days during which animals were allowed to move freely in the radial maze. To complete the session, mice had to visit each arm until its end. Choice accuracy was measured as the percentage of correct responses for each pair.

#### Stage 1: Acquisition of 3-pair discriminations

The acquisition task consisted in learning the position of the food within the maze. Indeed, each animal was assigned three adjacent pairs of arms (pairs A, B and C). In each pair, only one arm was baited as a food reward. The experiment is designed in a way that the left arms of pairs A and B are baited while the right arm of pair C is. In each trial, mice were given access to a pair (either pairs A, B, or C). A choice was made when the subject reached the food well of an arm; this also closed the door of the non-chosen arm. The trial was finished as soon as the animal returned to the central platform. The subject was then confined to the central platform for 20s before the next discrimination trial began: this constituted the inter-pair interval (ITI). This interval was selected based on previous work showing that a 20-s ITI is sufficient to challenge temporal binding and reveal deficits in relational/declarative memory in this paradigm (Sellami et al., 2018).

Each daily session consisted of 20 consecutive trials comprising alternate presentations of pairs A, B, and C according to a pseudo-random sequence. A mouse was considered to reach criterion when its overall choice accuracy was at or above 75% over two consecutive sessions given that performance in each of the three discrimination choices were at least 62% correct. The flexibility test task began the following day when criterion performance was reached. Lateralization index was quantified by using the absolute value of the subtraction of the performance on the left-baited pairs vs. the right-baited pairs. Lat = |((Pair A + Pair B) /2) - Pair C|. We then used the maximum lateralization over 3 consecutive days.

#### Stage 2: Flexibility probe of DM

In the test task, the position of the food in the maze did not change, but characteristic flexibility of DM expression was assessed by changing the way of presenting the arms. Indeed, in place of the pairs A and B, a pair AB was submitted to the mouse. The AB pair consisted in the combination of the two adjacent arms of the pairs A and B. It was the critical test of flexibility. Two other pairs were used as control: the pair C that remained unchanged (“unchanged learnt control”, and pair N (=new) made of the two arms non-used in acquisition (“unlearnt control”).

### Fiber photometry

#### Surgery

Mice were injected unilaterally 4 weeks before behavioral experiments with an Adeno-Associated Virus (AAV) GCaMP6f (AAV1-hsyn-jGCaMP6f-WPRE, Addgene). We used glass pipettes (tip diameter 25-35 µm) connected to a picospritzer (Parker Hannifin Corporation) into the right side of the dorsal CA1 of the hippocampus (0.1 µl/site; AP -1.8 mm; L +1.35 mm; DV -1.4 mm). Mice were then implanted with unilateral optic fiber implant (diameter: 200µm; numerical aperture: 0.39; flat tip; Neurophotometrics). Implants were fixed to the skull with Super-Bond dental cement (Sun Medical, Shiga, Japan). Mice were perfused after experiments to confirm correct placements of fibers. Viral injections targeted the pyramidal cell layer of the dCA1.

#### Recordings

Data were recorded throughout the entire fear conditioning paradigmusing a Neurophotometrics FP3002 system (Neurophotometrics, San Diego, CA). Briefly, 470 nm and 415 nm LED light was bandpass filtered, reflected by a dichroic mirror, and focused onto a multi-branch patch cord (Neurophotometrics, San Diego, CA) by a x20 objective lens. Alternating pulses of 470 and 415 nm light (∼50 µW) were delivered at 60 Hz, and photometry signals were time-locked to behavior using a Bonsai workflow. Signals were analyzed using the Synaptech suite cloud service (MetaCell) and custom-made Python scripts (Code available at https://github.com/elisecr/fiberphotometry_2024). Through the Synaptech platform, 470 and 415 signals were deinterleaved. For all experiments, the exponential delay period (the first 10 seconds(Hon et al., 2025)) of the recording was removed. To correct for photobleaching, isosbestic (415nm) signals were fit to a biexponential curve to model the baseline signal with photobleaching. The fitted biexponential curve was then scaled to match the 470nm signal using non-negative robust linear regress and subtracted from the 470 trace to correct for baseline drift. Traces were then Z-scored.

#### Experimental traces

Z scored data were analyzed by custom python scripts to extract calcium fluorescence traces for each experiment. For the trace fear conditioning experiments, these data were aligned to files generated by the poly fear software (Imetronic, France) “.dat” files to align the beginning of the tone and shock events to the closest millisecond in the Z scored data. The timing of the tones and trace periods were extracted as described above to accurately align the data.

#### Peak finding

The threshold for peak analysis definition was set at 1.0. Peak analysis was conducted using a custom-built Python script to calculate peak frequency and amplitude. Code available at https://github.com/elisecr/fiberphotometry-analysis

### Optogenetic Manipulation of dCA1

#### Surgery

Mice were injected bilaterally 5 weeks prior to behavioral experiments with an Adeno-Associated Virus (AAV) containing the excitatory opsin channelrhodopsin targeting the pyramidal cells of dCA1 (pAAV-CaMKIIa-hChR2(H134R)-mCherry (#26975), Addgene). We used glass pipettes (tip diameter 25-35 µm) connected to a picospritzer (Parker Hannifin Corporation) into the dorsal CA1 of the hippocampus (0.2 µl/site; AP -1.8 mm; L +1.3 mm; DV -1.4 mm and AP -2.5 mm; L +2 mm; DV -1.4 mm). Mice were then implanted with bilateral optic fiber implant (AP -1.8 mm; L +1.3 mm; DV -1.3 mm; diameter: 200µm; numerical aperture: 0.39; flat tip; Neurophotometrics). Implants were fixed to the skull with Super-Bond dental cement (Sun Medical, Shiga, Japan). Mice were perfused after experiments to confirm correct placements of fibers. Viral injections targeted the pyramidal cell layer of the dCA1. *Optogenetic manipulation during trace fear conditioning:* Mice were conditioned under a 40s trace interval condition, while dCA1 pyramidal neurons were activated <u>during the 40 seconds</u> <u>of trace interval only</u>. The light (≈3 mW per implanted fiber) was bilaterally conducted from the laser (Neurophotometrics, 450nm; 5 Hz; 5 ms laser on, 195 ms laser off) to the mice via two fiber-optic patch cords (diameter, 200 μm; Doric Lenses). The tone test was performed 24h later, mice were tethered to the patch chord, without optogenetic manipulation.

##### Open field

Mice were placed in a 38.5cm diameter circular arena and allowed free exploration for 5 minutes. These movements were recorded, and the mean velocity, distance travelled, and resting time were tracked and quantified using a custom MATLAB (Mathworks) code as in (Parkinson et al., 2024).

##### Immunohistochemistry

Naïve (homecage) and trained (90 min after the flexibility test on Day 3 for c-Fos analysis) animals were perfused transcardially with 0.01M phosphate buffered saline (PBS) to eliminate blood and extraneous material, followed by 4% paraformaldehyde (PFA). Brains were postfixed for 36h in PFA. Tissues were sectioned at 40 µm using a Leica 1000S vibratome and kept in a cryoprotective ethylene glycol solution at -20°C until processed for immunofluorescence. Sections were first washed and permeabilized in PBS-Triton 0.25%, then non-specific binding sites were blocked by immersing the tissue in 10% normal donkey serum, 2% bovine serum albumin in PBS-Triton 0.25% during 2h. Tissues were then stained using the primary antibodies overnight: mouse anti-c-Fos (1:1000; Santa Cruz). After 3x 15 min washes, we added anti-rabbit Alexa 488 (1:200; Life Technologies) secondary antibodies for 2h. After 3x 15 min washes slices were stained during 5 min with DAPI (5µM; Sigma), mounted on Livingstone slides then covered with Mowiol (Sigma) and coverslip (Thermofisher). c-Fos, staining from 2-3 slices par animals were imaged using a Nikon Axioscan confocal fluorescent microscope (20x objective). Stained sections of WT and mutant mice were imaged during the same imaging session. Immunofluorescence signals were quantified using the quPath software with routine particle analysis procedures (size= 30-500; circularity=0.30-1.00), to obtain cellular masks, divided by the area to obtain cell density per mm^2^.

##### Statistics

Data are presented as mean ± SEM. Statistical analyses were performed using GraphPad prism or StatView software for 2-way ANOVA, followed by Bonferroni *post-hoc* test to assess 2 variables or more (e.g., interaction between genotype/group and performance/freezing/number of cFos+ neurons) or Student’s t test, when assessing one variable only. Normality of the data was confirmed using the Kolmogorov–Smirnov test. Statistical significance was considered at p<0.05.

## Data availability

Raw data is provided in Supplementary Information S3. Code available at https://github.com/elisecr/fiberphotometry_2024

## Supporting information

Raw Data

## Acknowledgments

We thank all the personnel of The Australian Phenomics Facility and of The Neurocentre Magendie involved in mouse care.

## Author contributions

Conceptualization: ASA;

Methodology: ASA, ND, AM, MHM;

Investigation: ASA, ECR

Writing—original draft: ASA, ND

Writing—review & editing: ND, ASA, AM, ECR, MHM

## Supporting information

**Supplementary Figure S1:**
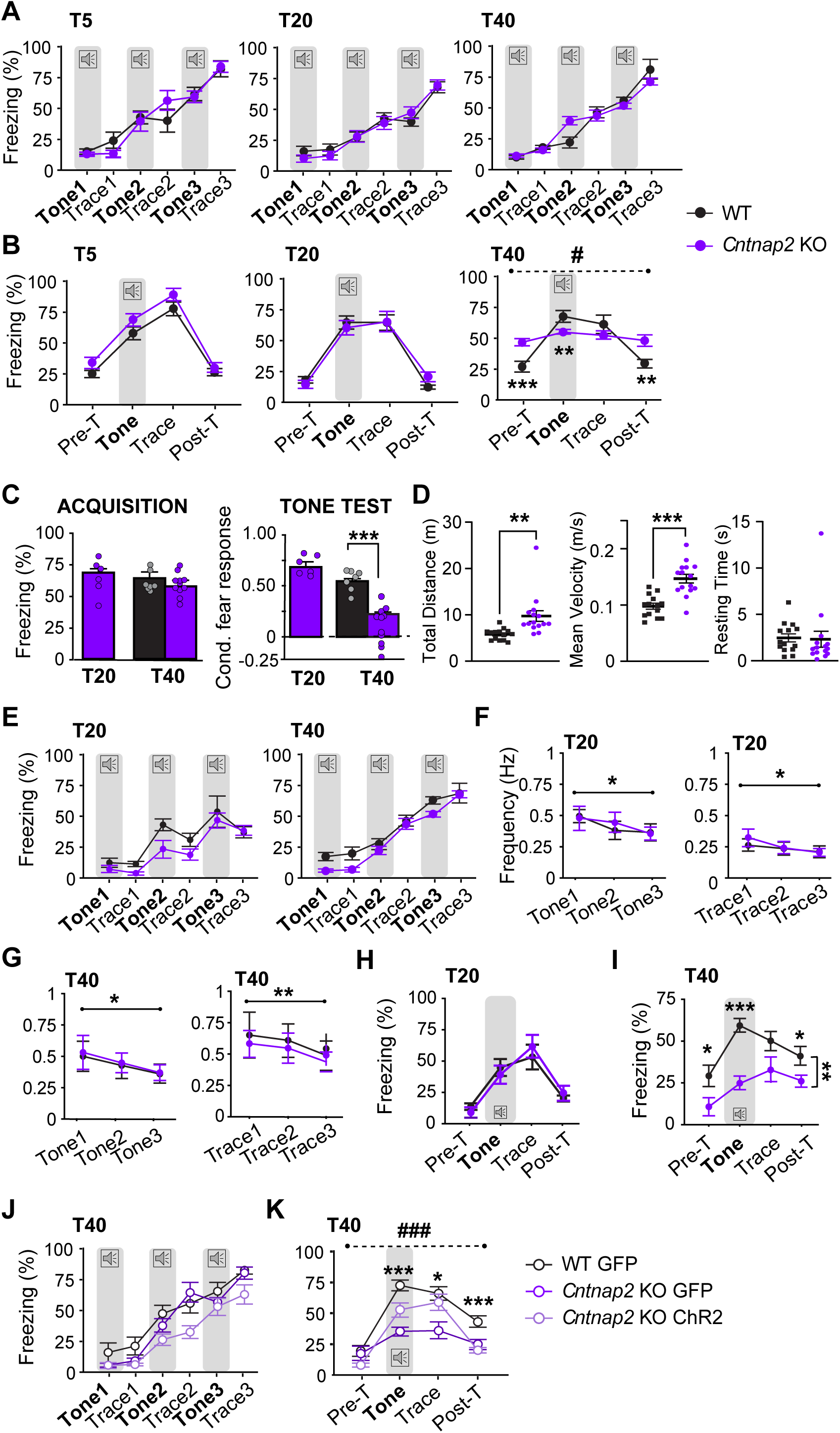
Freezing levels and calcium peak frequencies during the fiber photometry recordings (Figure 1). **A.** Freezing levels (percent) during the conditioning session for WT (black) and *Cntnap2* KO (purple) groups, conditioned with distinct trace durations (T5: 5s trace interval, T20: 20s trace interval; T40: 40s Trace interval). **B.** Freezing levels (percent time freezing) during the pre-tone, tone, trace and post-trace periods in the tone test for WT (black) and *Cntnap2* KO (purple) groups, 24h post-conditioning with distinct trace durations (T40: Pre-tone: p=0.001; Tone: p=0.0177; Trace: p=0.261; Post-Tone: p=0.0064; Genotype x Freezing interaction: F_3, 72_=12.671; p<0.0001). **C. Left:** Freezing levels (percent) during the third tone of the conditioning session for littermate WT (black) and *Cntnap2* KO (purple) T20 and T40 groups, (T20: *Cntnap2* KO n=6; T40: WT n=8; *Cntnap2* KO n=12). **Right:** Conditioned fear response during the tone test for the littermate groups (i.e., ratio of freezing before the tone compared to during the tone and the trace, see methods), 24h post-conditioning with distinct trace durations (Trace in *Cntnap2* KO: p<0.0001; Genotype in T40: p<0.0001). **D.** Analysis of locomotion in an open field. **Left:** Total distance (m); genotype: p=0.046. **Middle:** Mean velocity (m/s); genotype: p<0.0001. **Right:** Resting time (s); genotype: ns; n= 14 WT, 15 *Cntnap2* KO. **E. Fiber photometry**: Freezing levels (percent) during acquisition of trace fear conditioning for WT (black) and *Cntnap2* KO (purple) groups, conditioned with a 20s (T20, n=8 WT and n=8 *Cntnap2* KO, left panel) or a 40s trace interval (T40, n=11 WT and n=10 *Cntnap2* KO, right panel). **F. Left:** Peak frequency of Ca^2+^ transients during the 3 <u>tone presentations</u> of the T20 conditioning (tone effect: p=0.0176). **Right:** peak frequency of Ca^2+^ transients during the 3 <u>trace intervals</u> of the T20 conditioning (trace effect: p=0.0287). **G. Left:** peak frequency of Ca^2+^ transients during the 3 <u>tone presentations</u> of the T40 conditioning (tone effect: p=0.0018)**. Right:** peak frequency of Ca^2+^ transients during the 3 <u>trace intervals</u> of the T40 conditioning (trace effect: p=0.0229). **H.** Freezing levels (percent) during the tone test of the fiber photometry (T20) for WT (black) and *Cntnap2* KO (purple) groups, 24h post-T20 conditioning (ns; WT n=8; KO n=8). **I.** Freezing levels (percent) during the tone test of the fiber photometry (T40) for WT (black) and *Cntnap2* KO (purple) groups, 24h post-conditioning (Genotype: p=0.0009; PreTone: p=0.0258; Tone: p<0.0001; Post-Tone: p=0.0177; WT n=11; KO n=10). **J. Optogenetic**: Freezing levels (percent) during acquisition for GFP-injected WT (black; n=6), GFP-injected *Cntnap2* KO (purple; n=7), and ChR2-injected *Cntnap2* KO (light purple, white circles n=11) groups, conditioned with a 40s trace interval (Group: ns). **K.** Freezing levels (percent) during the tone test for GFP-injected WT (black; n=6), GFP-injected *Cntnap2* KO (purple; n=7) and ChR2-injected *Cntnap2* KO (light purple white circles n=11) groups, 24h post-T40 conditioning (Group x freezing: F_6, 63_ = 5.656; p<0.0001; Tone: p=0.0009; Trace: p=0.0224; Post-T: 0.0004).

**Supplementary Figure S2:**
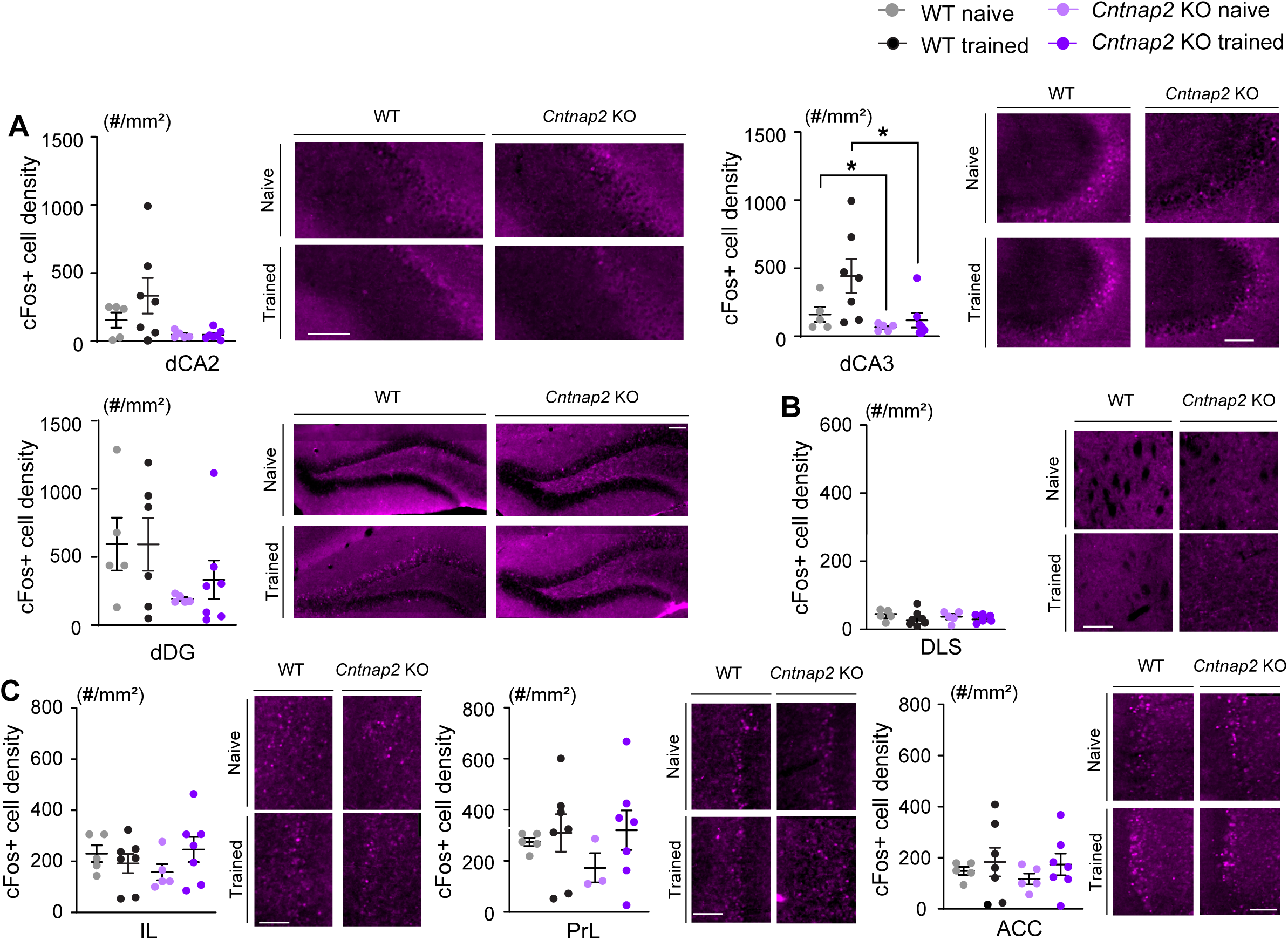
cFos density in the hippocampus, striatum, and prefrontal cortex. **A.** Number of cFos-positive neurons and representative images (cell density/mm^2^, scale: 100µm) in the dorsal hippocampus subfields: dCA2, dCA3 (Naive WT vs. KO: p=0.0400; Trained WT vs. KO: p=0.0454), dDG for naïve (light colors) and trained (dark colors) WT (grey/black) and *Cntnap2* KO (purple) groups (WT Naïve: n=5; WT Trained: n=7; *Cntnap2* KO: n=5; *Cntnap2* KO Trained: n=7). **B.** Number of cFos-positive neurons (cell density/mm^2^) in the Dorso-Lateral Striatum (DLS; ns). **C.** Number of cFos-positive neurons (cell density/mm^2^) in the prefrontal cortex: Infra-Limbic, Pre-Limbic, Anterior Cingular Cortex (PFC, IL, PrL, ACC, respectively). Data shown at mean ± SEM. *: p<0.05.

